# White-matter connectivity shapes visual and semantic representations in the aging brain

**DOI:** 10.64898/2026.04.07.716890

**Authors:** Cortney M. Howard, Shenyang Huang, Kirsten Gillette, Lifu Deng, Roberto Cabeza, Simon W. Davis

**Author notes:** Corresponding Author: Cortney Howard Duke University Durham, NC 27708. The authors declare no conflicts of interest.

## Abstract

Multiple lines of evidence indicate that aging is associated with a reduction in the quality of sensory information coupled with increased reliance on semantic information. Consistent with this evidence, functional MRI studies have repeatedly found that, compared to younger adults, older adults show weaker visual representations but stronger semantic representations. However, the factors that engender this visuo-semantic representational shift (VSRS) remain unclear. Here, we test whether structural connectivity within the representational network predicts these shifts. In the face of typical age-related declines in visual memory and structural connectivity, we find that structural connectivity (i) mediates age-related decline in perceptual representation, and (ii) supports semantic and perceptual mnemonic representations in older adults. These findings outline a potential mechanism of age-related perceptual processing declines and highlight the adaptability of the memory system with age.

## Main Text

A central challenge in cognitive aging is to explain why some neural systems and cognitive functions are especially vulnerable to age-related decline, whereas others remain relatively resilient and continue to support performance despite that decline. In this framework, *vulnerability* refers to susceptibility to age-related declines in neural integrity or cognitive performance, whereas *resilience* refers to the capacity to maintain or support performance despite such declines (Bocancea et al., 2021; Stern et al., 2023). In the domain of object cognition, converging behavioral and neuroimaging evidence indicates that aging disproportionately compromises the processing of visual features (contrast sensitivity; motion, color, and shape perception, etc.; for a review, see Owsley, 2016) and memory for perceptual detail (Delarazan et al., 2023; Monge & Madden, 2016; Naveh-Benjamin & Kilb, 2014), while semantic knowledge and semantic processing are often preserved or even strengthened (Spreng & Turner, 2019; Umanath & Marsh, 2014). Recent fMRI studies point to a parallel pattern at the representational level in occipito-temporal cortex: compared with younger adults, older adults show weaker visual representations for objects in visual representation regions (Bowman et al., 2019; Hill et al., 2021; Koen et al., 2020; Srokova et al., 2024), alongside stronger semantic representations for the same stimuli in semantic representation regions (Deng et al., 2021; Naspi et al., 2023; Pauley & Sander, 2025). We refer to this pattern as a visuo-semantic representational shift (VSRS). Together, these findings suggest that age-related memory change may reflect not only vulnerability in sensory systems, but also resilience in semantic systems. The neural architecture that gives rise to this shift, however, remains unknown.

A key unanswered question is whether the VSRS is constrained by the structural architecture of the representational network. This possibility is particularly timely in light of increasing interest in visual system white matter in the neuroscience of aging (Alam et al., 2024; Liu et al., 2025; Rokem et al., 2017). In cognitive aging, white matter pathways linking visual representation and semantic regions may be especially important for two reasons. First, age-related decline in these pathways could exacerbate vulnerability by weakening the integrity of visual representations. Second, preserved connectivity within the same network could enable resilience by supporting access to semantic representations when visual representations are degraded. Although structural decline in ventral visual pathways has been linked to age-related changes in perception and memory (Raz & Rodrigue, 2006; Rieck et al., 2020), the relationship between white matter integrity and representational change itself remains poorly understood. Addressing this uncertainty is critical, as recent work suggests that age-related changes in neural representations are pervasive across encoding, retrieval, and reinstatement. Thus, the present study addresses this gap in knowledge by asking whether structural connectivity helps determine where such representational vulnerability emerges and when more resilient semantic mechanisms can be recruited. Age-related degradation in visual representation has been linked directly to white matter microstructure (Rieck et al., 2020) and OTC decline has been associated with deficits in both visual working memory and episodic memory (Bennett & Rypma, 2013; Davis et al., 2009, 2018; Duarte et al., 2010; Lockhart et al., 2012). In the visual domain, studies by Behrman (Thomas et al., 2008) and Saygin (Saygin et al., 2012) have found that reductions in white matter connections in the OTC network may account for age-related changes in face perception. This evidence has led some to suppose that visual perception deficits cascade through the cognitive system, disrupting memory and executive functions (Monge & Madden, 2016). Thus, structural decline in the OTC plays a crucial role in age-related cognitive decline.

Beyond the occipito-temporal cortex, inferior parietal cortex (IPC) may also contribute to semantic resilience in aging. IPC has been implicated in semantic integration (Bonner et al., 2013; Kuhnke et al., 2023; Price et al., 2016), and in the transformation of representations across encoding and retrieval (Favila et al., 2018; Howard et al., 2024). Given that IPC is strongly linked to semantic processing, representations may be more visual/concrete during encoding but more semantic/abstract during retrieval, which implicates the IPC as a semantic and integration hub (Favila et al., 2020) in younger adults. If so, connectivity between IPC and occipito-temporal regions may support the transformation of perceptual representations into more abstract semantic representations in older adults. Critically, this raises the possibility that resilience does not depend solely on temporal-lobe semantic regions, but on a broader network linking visual representation regions with temporal and parietal areas that support semantic abstraction and memory-guided behavior. Clarifying how age-related changes in the mnemonic representations affected by VSRS are constrained by structural architecture is critical for understanding whether such shifts represent a strictly compensatory mechanism or a fundamentally new functional strategy of encoding real-world stimuli.

The current study addresses two related questions. First, does age-related decline in structural connectivity contribute to weaker visual representations in the representational network, thereby linking structural vulnerability to functional vulnerability? Second, can preserved semantic representations help support memory performance in older adults despite declines in visual processing? To address these questions, we combined fMRI-based RSA during object perception, diffusion-weighted imaging of structural connectivity within the representational network, and behavioral measures from conceptual and perceptual memory tests. We hypothesized that older adults would show lower perceptual memory, weaker structural connectivity, and weaker visual representations. We further asked whether preserved semantic representations might provide a source of resilience, particularly under conditions of reduced visual-system integrity.

## Results

### Perceptual memory performance

To examine whether perceptual memory was impaired in older adults (OAs) compared to younger adults (YAs), we performed two-sample t-tests on their corrected recognition (CR). Older adults had significantly lower CR scores than YAs on the perceptual memory test. In contrast, age groups did not differ significantly on conceptual memory, consistent with evidence that aging disproportionately affects memory for perceptual detail (Delarazan et al., 2023; Monge & Madden, 2016) (**Figure 1D**, **Table 1**).

**Table 1.** Age Effects in Corrected Recognition.

|  | Mean YA | SE YA | Mean OA | SE OA | t | p |
| --- | --- | --- | --- | --- | --- | --- |
| Perceptual CR | 0.498 | 0.042 | 0.365 | 0.034 | 2.455 | 0.018 |
| Conceptual CR | 0.532 | 0.031 | 0.493 | 0.041 | 0.762 | 0.450 |

\*Note: CR – corrected recognition, SE – standard error, YA – younger adults, OA – older adults

### Representational strength

To examine the degree to which our sample exhibited age-related visuo-semantic representational shifts (VSRS), tRSA values were analyzed using a series of mixed-effects linear models. For each broad region of interest, we constructed a model that included the interaction between Age Group (older adults and younger adults) and Representation Type (semantic and visual), their main effects, and random intercepts for Participant, Stimuli, and Sub-Region of Interest.

We classified these regions based on the nature of the Age Group by Representation Type interaction. The lateral occipital cortex (LOC), fusiform gyrus (FUG), parahippocampal gyrus (PHG), anterior inferior temporal cortex (aITC), and anterior inferior parietal cortex (aIPC) exhibited significant Age Group by Representation Type interactions. These interactions are driven by different patterns across regions. In LOC and FUG, older adults exhibited a greater deficit in visual representations than in semantic representations. In PHG and aITC, in contrast, the interaction reflected opposite effects of aging on visual and semantic representations: compared to young adults, older adults showed weaker visual representations but stronger semantic representations. Stronger semantic representations in older than younger adults were also found in aIPC, although without a significant difference in visual representations. Lastly, EVC, pITC, and pIPC displayed significant visual and/or semantic representations but did not show reliable Age Group: Representation Type interactions (**Figure 3**, **Table 2**).

**Figure 1:**
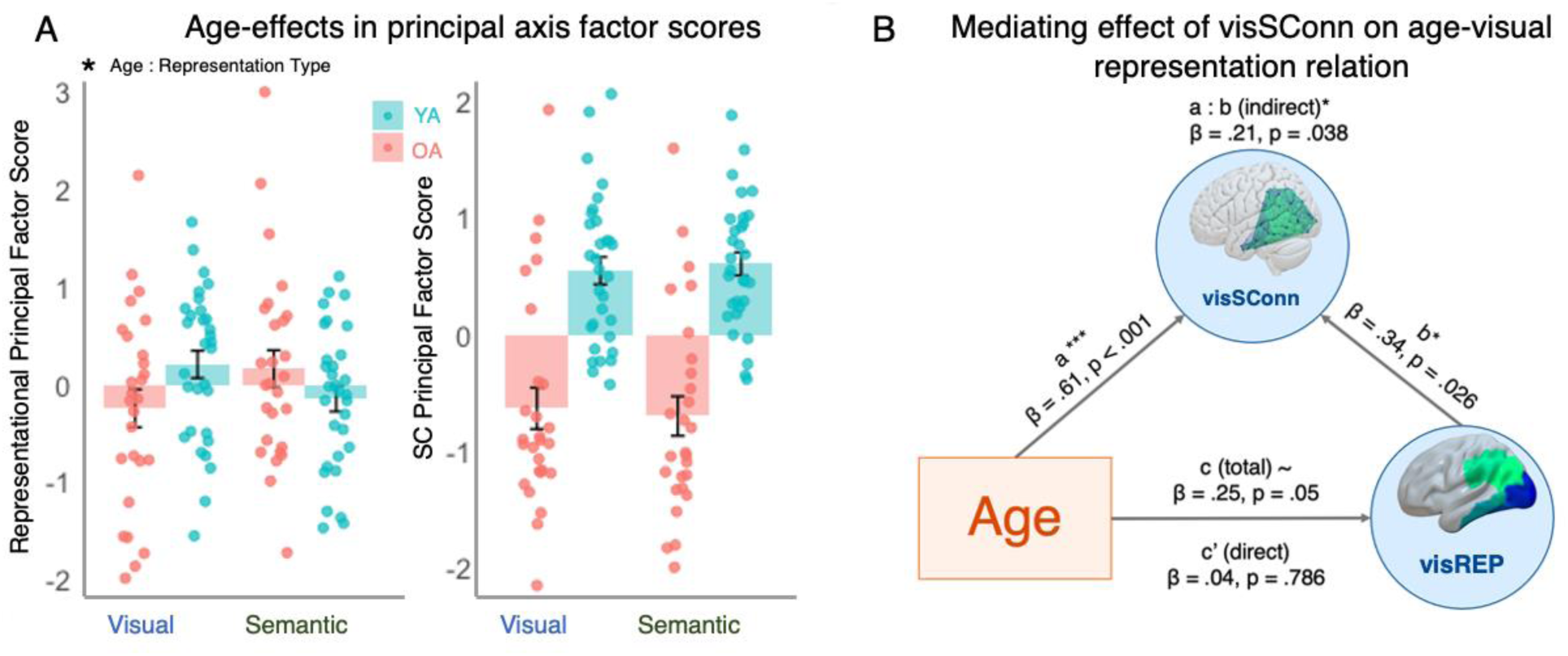
Structural Connectivity Mediates the Age – VSRS Relationship. *A) Age-effects in principal axis factor scores*. The representational principal factor scores (left, y axis) exhibit a significant age (YA in teal, OA in salmon) by representation type interaction such that visual representations (blue) show an age-related decline while semantic representations (green) show an age-related boost. The structural connectivity (SC) principal factor scores (right, y-axis) exhibit significant age-related decline for both visual (blue) and semantic (green) representation regions. *B) Structural connectivity in visual representation regions (visSConn).* visSConn significantly mediates the age-visual representation relationship.

**Figure 3:**
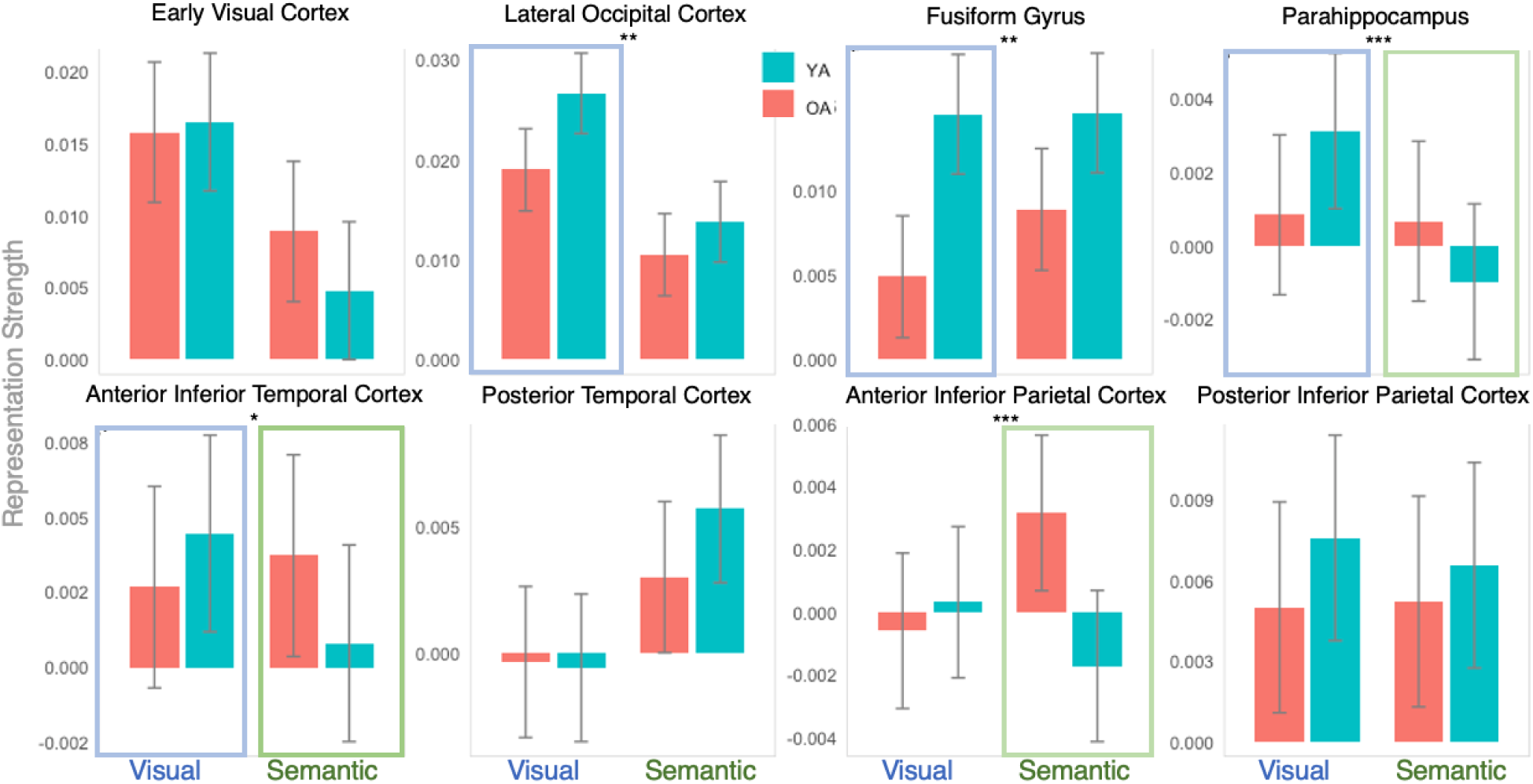
Age-Related Visuo-Semantic Representational Shifts (VSRS). Results from the mixed-effects model detecting VSRS regions. Four broad regions exhibited age-related declines in visual (blue) representation: lateral occipital cortex, fusiform gyrus, parahippocampus, and anterior inferior temporal cortex (blue outline). Three broad regions of interest exhibited age-related increases in semantic (green) representation: parahippocampus, anterior inferior temporal cortex, anterior inferior parietal cortex (green outline). Data from older adults (OA) are shown in salmon bars while data from younger adults (YA) are shown in teal. Significant Age by Representation Type interactions are indicated by black asterisks. FDR corrected * p < 0.05; ** p < 0.01, *** p < 0.005.

**Table 2.** Age: Representation Type Interactions.

| ROI | DenDF | F | p | FDR adjusted |
| --- | --- | --- | --- | --- |
| EVC | 27938 | 2.599 | 0.107 | 0.143 |
| LOC | 84173 | 6.713 | 0.010 | 0.026 |
| FUG | 84173 | 5.727 | 0.017 | 0.027 |
| PHG | 167745 | 12.135 | <.001 | 0.002 |
| aITC | 55868 | 5.818 | 0.016 | 0.027 |
| pITC | 56055 | 2.236 | 0.135 | 0.154 |
| aIPC | 112291 | 17.681 | <.001 | <0.001 |
| pIPC | 56055 | 0.392 | 0.531 | 0.531 |

\*Note: ROI – region of interest, EVC – early visual cortex, LOC – lateral occipital cortex, FUG – fusiform gyrus, PHG – parahippocampus, aITC – anterior inferior temporal cortex, pITC – posterior inferior temporal cortex, aIPC – anterior inferior parietal cortex, pIPC – posterior inferior parietal cortex, SE – standard error, FDR adj – false discovery rate adjustment

Overall, these findings are consistent with an age-related visuo-semantic representational shift (VSRS). Specifically, compared to younger adults, older adults exhibited weaker visual representations than young adults in LOC, FUG, PHG, and aITC, and stronger semantic representations in PHG, aITC, and aIPC.

### Structural Connectivity

We utilized the DWI data from the high-resolution MUSE scan to construct structural connectomes to investigate the structural connectivity of the eight representation regions. This connectome captures both the number of streamlines connecting each pair of regions and the average fractional anisotropy (FA) within those streamlines. For each representation region of interest (ROI), we calculated the average connection strength to the other representation regions for each participant in the study. This connection strength was then analyzed using a two-sample t-test to determine whether it was significantly greater in younger adults (YA) compared to older adults (OA). Our findings revealed that, across all broad regions, the structural connection strength was indeed greater in YA than in OA, confirming the age-related decline in structural connectivity in representation regions (**Figure 4a**, **Table 3**).

**Figure 4:**
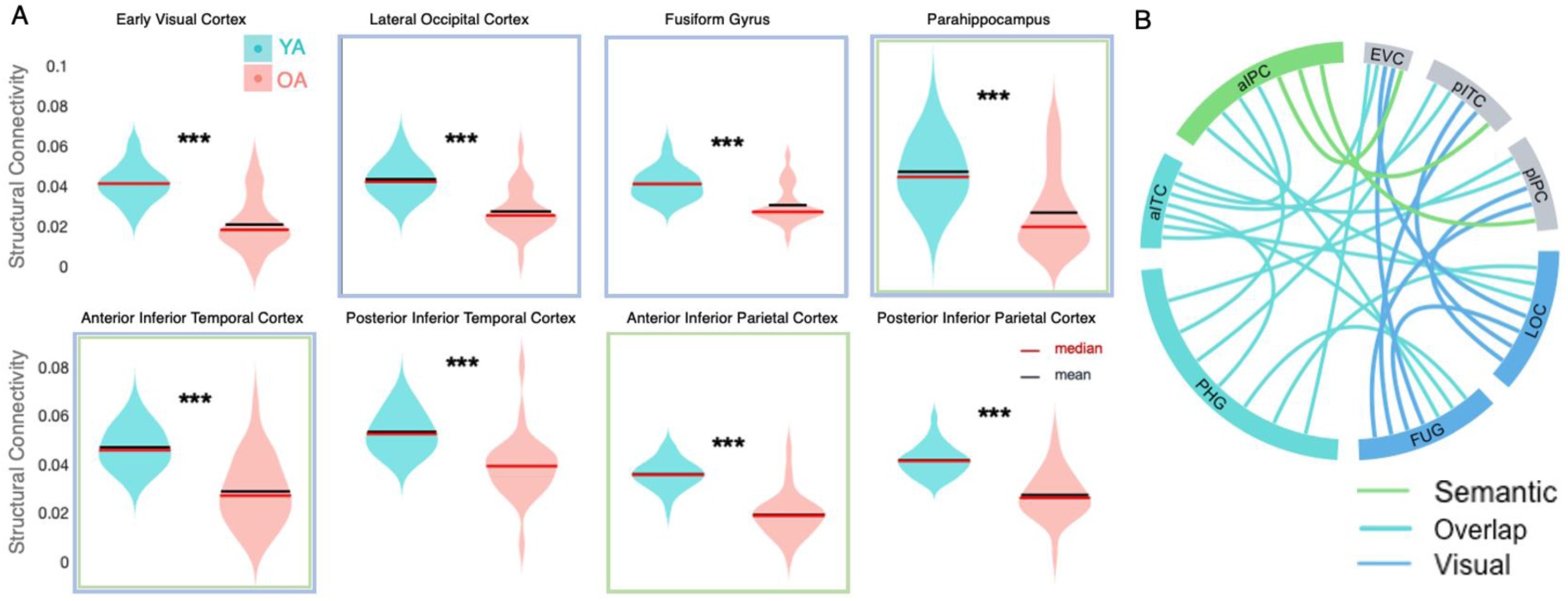
Age – Related Declines in Structural Connectivity Among Representation Regions. **A**) *Structural connectivity among representation regions.* All representation regions show robust age-related declines in structural connectivity (red line: median; black line: mean). Regions showing age-related declines in visual representation are outlined in blue, and regions displaying age-related increases in semantic representation, in green (see Figure 3, *** p < 0.005. **B**) *Connections considered Semantic (green) or Visual (blue), and their overlap (teal)*.

**Table 3.** Age Effects in Structural Connectivity.

| ROI | Mean YA | SE YA | Mean OA | SE OA | t | p | FDR adj |
| --- | --- | --- | --- | --- | --- | --- | --- |
| EVC | 0.041 | 0.002 | 0.021 | 0.003 | 6.868 | < 0.0001 | < 0.0001 |
| LOC | 0.043 | 0.002 | 0.027 | 0.002 | 6.134 | < 0.0001 | < 0.0001 |
| FUG | 0.041 | 0.001 | 0.030 | 0.002 | 4.850 | < 0.0001 | < 0.0001 |
| PHG | 0.047 | 0.003 | 0.027 | 0.004 | 4.368 | 0.0001 | 0.0001 |
| aITC | 0.047 | 0.002 | 0.029 | 0.003 | 6.175 | < 0.0001 | < 0.0001 |
| pITC | 0.053 | 0.001 | 0.039 | 0.002 | 5.042 | < 0.0001 | < 0.0001 |
| aIPC | 0.036 | 0.001 | 0.019 | 0.002 | 8.490 | < 0.0001 | < 0.0001 |
| pIPC | 0.042 | 0.001 | 0.027 | 0.002 | 6.700 | < 0.0001 | < 0.0001 |

\*Note: ROI – region of interest, YA – younger adults, OA – older adults, EVC – early visual cortex, LOC – lateral occipital cortex, FUG – fusiform gyrus; PHG – parahippocampus, aITC – anterior inferior temporal cortex, pITC – posterior inferior temporal cortex, aIPC – anterior inferior parietal cortex, pIPC – posterior inferior parietal cortex, SE – standard error, FDR adj – false discovery rate adjustment

### Mediation and moderation

Four *Principal Axis Factor* scores (PAFs) were generated using data from visual and semantic representational change regions, which were identified in a data-driven manner depending on the significant age-group effects in the representational mixed effects models. The first two PAFs corresponded to measures of visual and semantic representational strength and were generated using ROIs with significant or trend-level age-related differences in visual (LOC, FUG, PHG, and aITC) or semantic (PHG, aITC, and aIPC) representation strength, respectively. For these ROIs, the model estimate of visual or semantic representational strength for each participant was extracted from the representational mixed effect model described above. The third and fourth PAFs corresponded to measures of structural connectivity from visual representation ROIs (LOC, FUG, PHG, and aITC) or semantic representation ROIs (PHG, aITC, and aIPC) to all other regions in the representation networks (**Table 4** lists all connections included in the structural connectivity calculations).

**Table 4.** Principal axis Factor Analyses.

| Measure | Visual Rep | Semantic Rep | Visual SC | Semantic SC |
| --- | --- | --- | --- | --- |
| Factor loadings (range) | 0.28–0.88 | 0.47–0.79 | 0.84–0.97 | 0.81–0.96 |
| Variance explained | 39% | 46% | 82% | 78% |
| Off-diagonal fit | 0.93 | 1.00 | 1.00 | 1.00 |
| Factor score<br>adequacy | $r = .91, R^2 = .84,$<br>min $r = .67$ | $r = .87, R^2 = .76,$ min<br>$r = .51$ | $r = .98, R^2 = .96,$ min<br>$r = .93$ | $r = .97, R^2 = .94,$ min<br>$r = .89$ |

\*Note: Rep – representation, SC – structural connectivity

Model adequacy was evaluated using indices of factor score adequacy, which assess how well the estimated factor scores represent the latent construct. Following conventional guidelines, correlations of factor scores with the latent factor above 0.70, multiple R² values above 0.50, and minimum factor score correlations above 0.50 are considered acceptable. Across all models, adequacy indices exceeded these thresholds (all r ≥ 0.87, R² ≥ 0.76, min r ≥ .51), supporting the use of single-factor scores for data simplification.

To ensure construct validity the visual and semantic representational strength factors were submitted to a linear model with the main effects of age group and representation type as well as their interaction. Consistent with the ROIs employed to generate visual and semantic representation PAFs, the linear models revealed a significant interaction of age * representation type because visual representations showed an age-related reduction whereas semantic representations displayed age-related increases. Thus, taken together, the representation PAFs reveal a clear VSRS and validate the PAFs measures (**Figure 5A left, Table 5**).

**Table 5.**
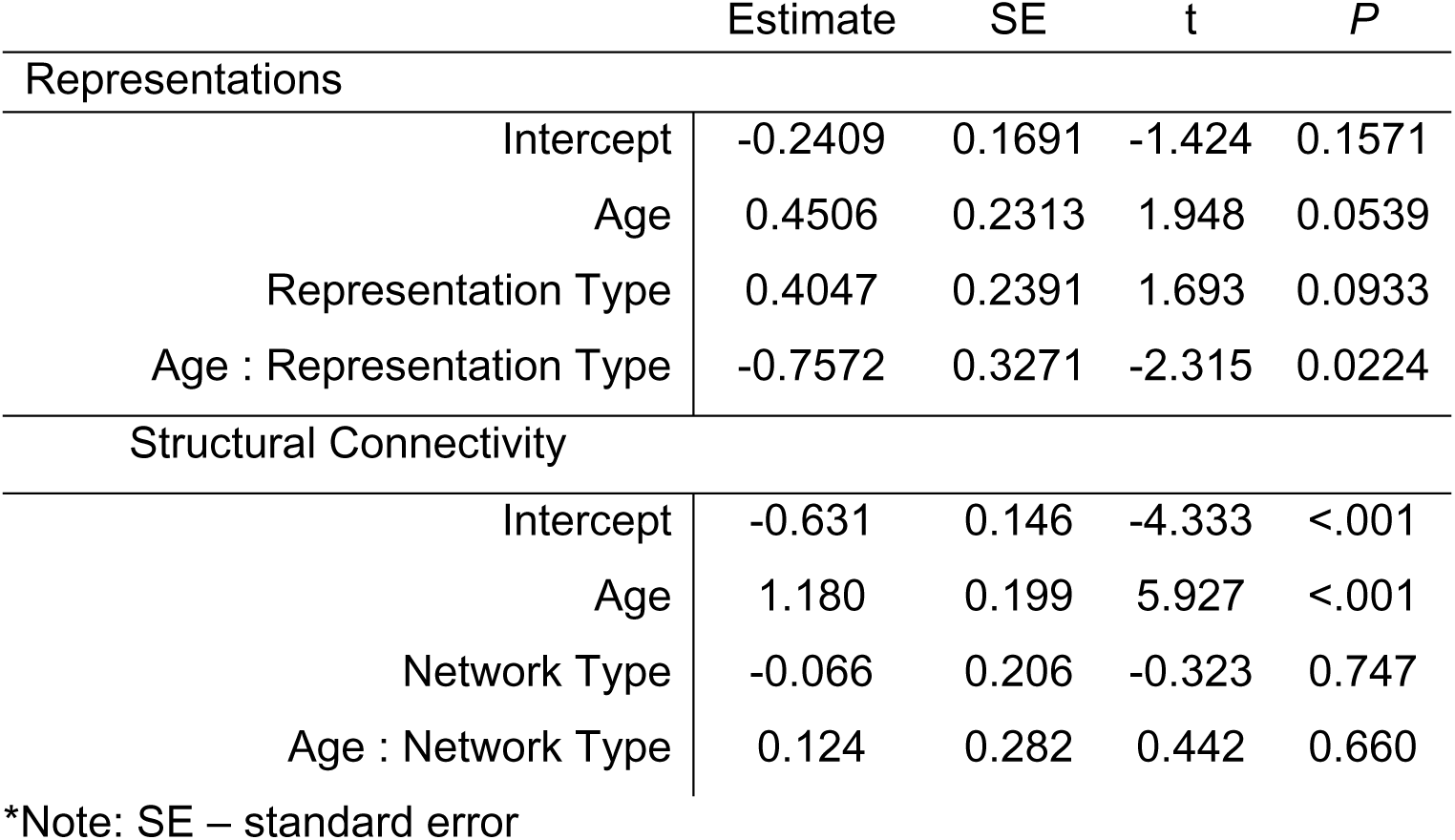
Linear Models of Principal axis Factors.

|  | Estimate | SE | t | P |
| --- | --- | --- | --- | --- |
| Representations |  |  |  |  |
| Intercept | -0.2409 | 0.1691 | -1.424 | 0.1571 |
| Age | 0.4506 | 0.2313 | 1.948 | 0.0539 |
| Representation Type | 0.4047 | 0.2391 | 1.693 | 0.0933 |
| Age : Representation Type | -0.7572 | 0.3271 | -2.315 | 0.0224 |
| Structural Connectivity |  |  |  |  |
| Intercept | -0.631 | 0.146 | -4.333 | <.001 |
| Age | 1.180 | 0.199 | 5.927 | <.001 |
| Network Type | -0.066 | 0.206 | -0.323 | 0.747 |
| Age : Network Type | 0.124 | 0.282 | 0.442 | 0.660 |

\*Note: SE – standard error

Similarly, the PAFs for structural connectivity of visual and semantic representation regions were consistent with the structural connectivity data, displaying a significant main effect of age group, but no interaction between age and region type. In other words, structural connectivity declined in OA relative to YA for both visual and semantic representation regions (**Figure 5A right, Table 5**).

Having confirmed the validity of representation strength and connectivity PAFs, we used mediation models to link the age-related deficit of visual representations (visREP) and the age-related decline on the structural connectivity of visual representation regions (visSConn). The model revealed significant effects of age group on visSConn (a path), visSConn on visREP (b path), and mediation (indirect effect, a*b) of visSConn on the relationship between age and visREP. The Age group on visREP (total, c path) was trending (Z = 1.95, p = 0.05), and the direct effect after controlling for visSConn was not significant (**Figure 5B**). This finding suggests that age-related declines in visual representation are accounted for by degraded structural connectivity between visual representation regions.

Having demonstrated the role of structural connectivity on age-related visual representation decline, we investigated the role of structural connectivity on semantic representations. Following the same logic, we investigated a mediation model linking the effects of age on semantic representations (semREP) and the age-related decline on the structural connectivity of semantic representation regions (semSConn). Unlike the visual mediation model, the semantic mediation model did not reveal any significant mediation effect. These results suggest that changes in structural connectivity can account for deficits in visual representations (**Table 6**) but not for enhanced semantic representations.

**Table 6.**
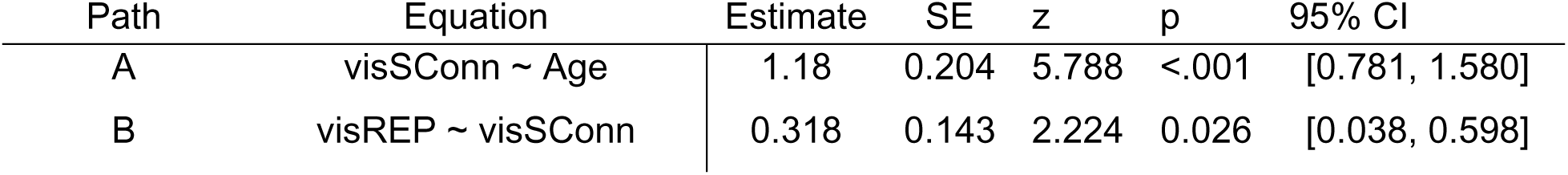
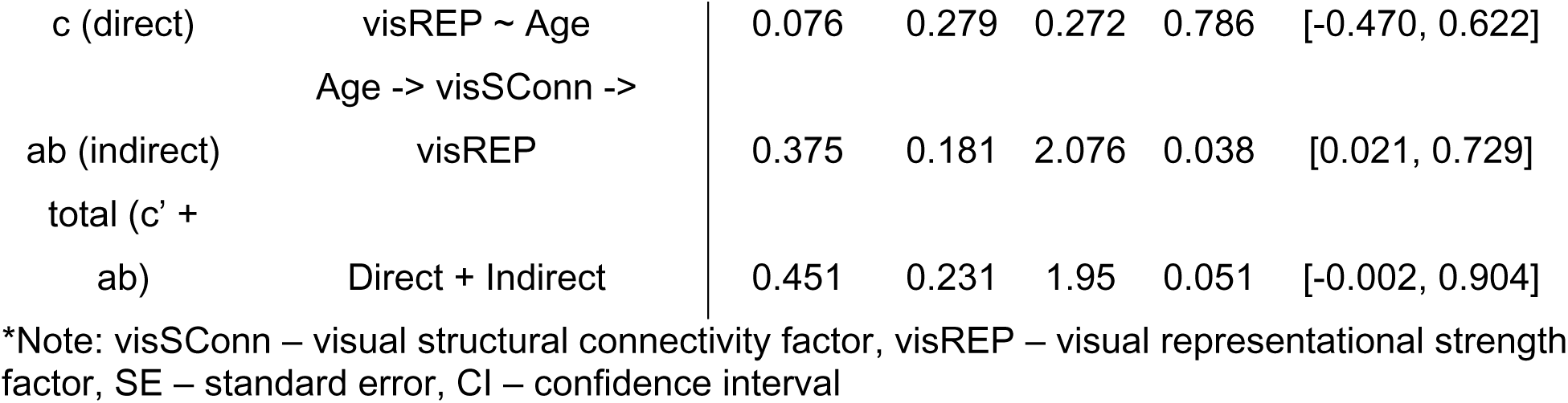
Mediation Results.

To further understand the dynamics of VSRS, moderation models assessed the interaction of age and structural connectivity on representation strength. The first model tested the extent to which visSConn moderates the relationship between age and visREP. The analysis revealed that visSConn positively predicts visREP (t = 2.722, p = 0.01). Additionally, a trending but not significant age by visSConn interaction (t = -1.6, p = 0.11) suggests visSConn positively predicts visREP most strongly in OAs (**Figure 6A**). This analysis tentatively confirmed our hypothesis that the impact of structural connectivity on visual representation strength only occurs later in life with sufficient degradation (**Table 7**).

**Figure 6:**
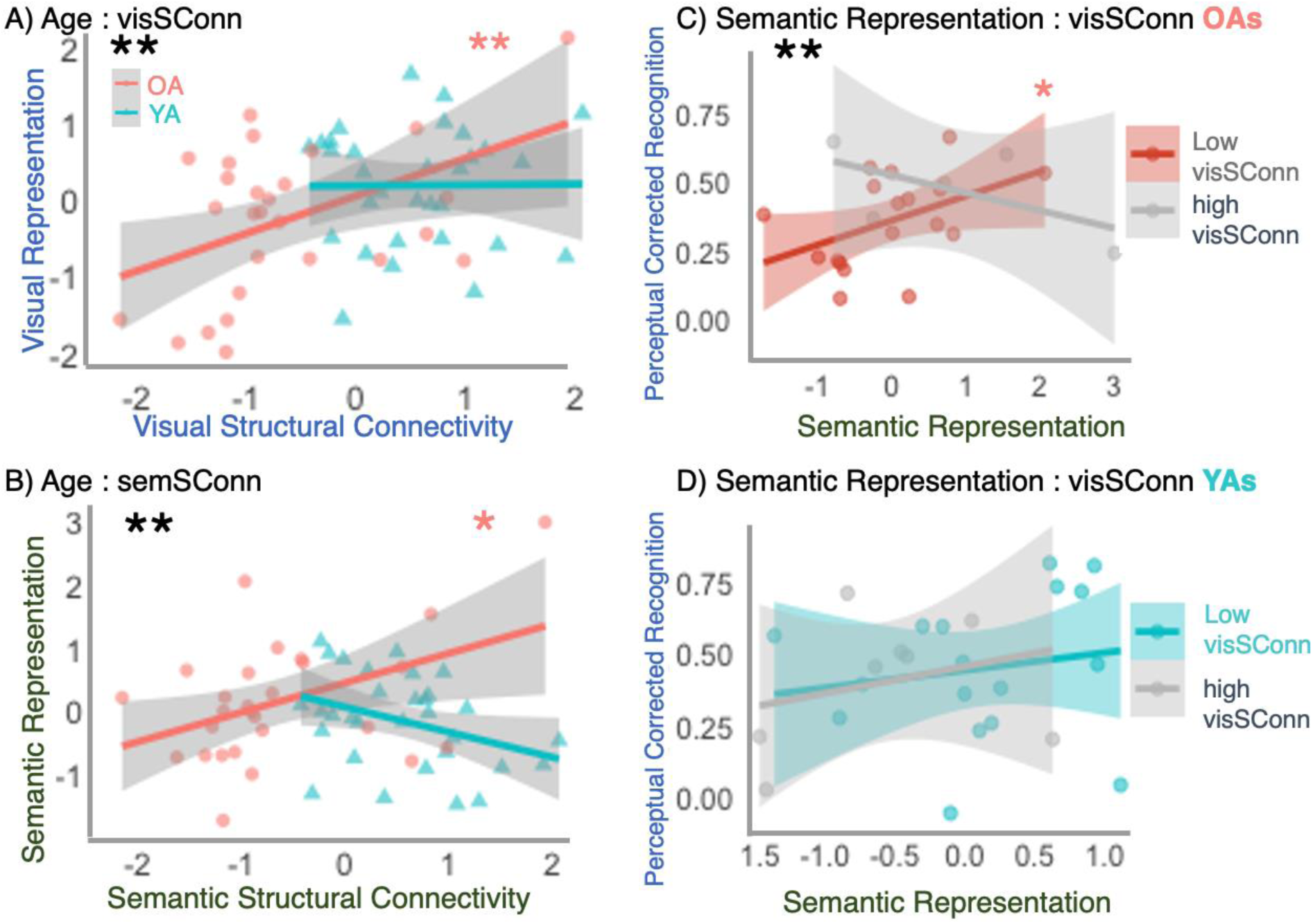
**Structural Connectivity Moderates VSRS and Older Adult Perceptual Memory Performance**. A) visSConn (x-axis) significantly moderates the age-visREP (y-axis) relationship such that older adult (salmon), not younger adult (teal) visSConn positively predicts visREP. B) semSConn (x-axis) significantly moderates the age-semREP (y-axis) relationship such that older adult (salmon), not younger adult (teal) semSConn positively predicts semREP. C) For older adult participants, visSConn significantly moderates the semREP (x-axis) - Perceptual Memory (y-axis) relationship such that only older adults with low visSConn exhibit a positive semREP – perceptual memory relationship. D) For younger adult participants, visSConn does not moderate the semREP (x-axis) – Perceptual Memory (y-axis) relationship.

**Table 7.** Moderation Results.

| Effect | Estimate | SE | T | p_value |
| --- | --- | --- | --- | --- |
| Age and semSConn interaction on semREP |  |  |  |  |
| Intercept | 0.490 | 0.202 | 2.425 | 0.019 |
| Age | -0.405 | 0.299 | -1.354 | 0.181 |
| semSConn | 0.468 | 0.182 | 2.570 | 0.013 |
| Age : semSConn | -0.843 | 0.326 | -2.585 | 0.013 |
| Age and visSConn interaction on visREP |  |  |  |  |
| Intercept | 0.071 | 0.200 | 0.354 | 0.724 |
| Age | 0.132 | 0.284 | 0.466 | 0.643 |
| visSConn | 0.494 | 0.182 | 2.722 | 0.009 |
| Age : visSConn | -0.483 | 0.300 | -1.608 | 0.114 |
| OAs: semREP and visSConn on perceptual memory performance |  |  |  |  |
| Intercept | 0.483 | 0.051 | 9.464 | <.001 |
| semREP | 0.007 | 0.036 | 0.196 | 0.847 |
| visSConn | 0.104 | 0.046 | 2.255 | 0.036 |
| semREP : visSConn | -0.075 | 0.026 | -2.934 | 0.009 |
| YAs: semREP and visSConn on perceptual memory performance |  |  |  |  |
| Intercept | 0.457 | 0.061 | 7.441 | <.001 |
| semREP | 0.058 | 0.090 | 0.639 | 0.530 |
| visSConn | 0.078 | 0.081 | 0.964 | 0.346 |
| semREP : visSConn | 0.054 | 0.114 | 0.474 | 0.640 |

\*Note: visSConn – visual structural connectivity factor, visREP – visual representational strength factor, semSConn – semantic structural connectivity factor, semREP – semantic representation factor, SE – standard error, CI – confidence interval

The second moderation model revealed similar findings for semSConn and semREP: for OAs but not for YAs, semSConn positively predicts semREP (**Figure 6B**), here the interaction is significant (t = -2.585, p = 0.01). This suggests that, while structural connectivity does not mediate the age-semantic representation relationship, older adults must have sufficiently intact connectivity of semantic representation regions to facilitate increased semantic representations (**Table 7**).

Next, we conducted moderation analyses separately for YAs and OAs to understand the effect of visSConn and semantic representation on perceptual memory performance. These analyses revealed that OAs (**Figure 6C**), and not YAs (**Figure 6D**), exhibit a significant interaction between visSConn and perceptual corrected recognition (t = -2.93, p = 0.01), such that only OAs with low visSConn showed a significant, positive relationship between semREP and perceptual corrected recognition. This result supports the hypothesis that the increase in semantic representation compensates for degraded structural connectivity within visual representation regions (**Table 7**).

## Discussion

This study examines how age-related changes in object memory reflect both vulnerability and resilience within the cortical representational system. Older adults showed reduced perceptual memory, weaker visual representations, and lower structural connectivity in visual representation regions, indicating vulnerability of visual and structural mechanisms. At the same time, older adults showed stronger semantic representations in semantic representation regions, and these semantic signals supported perceptual memory under conditions of reduced visual-system connectivity, indicating resilience. Critically, by integrating fMRI-based representational analyses with high-resolution diffusion imaging, the present study identifies structural connectivity as a key constraint on when age-related representational decline emerges and when semantic support can be recruited.

Our regional visual–semantic representational shift (VSRS) findings broadly converge with prior work showing that aging weakens visual object representations while preserving or enhancing semantic representations (Deng et al., 2021; Naspi et al., 2023). In the present study, age-related reductions in visual representational strength were most evident in LOC, fusiform, parahippocampal, and anterior inferior temporal regions. This pattern is consistent with the idea that age-related visual vulnerability is distributed across visual representation regions rather than confined to a single locus. Notably, a recent study examining age-related differences in representation using a model-free approach also found item-level, but not category-level, representations decline with age in the lateral occipital cortices (Srokova et al., 2024). This could be due to the semantic nature of category-level representations and underscores the importance of examining both category-level and item-level representational fidelity when interpreting VSRS phenomena. Importantly, these findings suggest that the cortical location of age-related representational changes may depend on the level of visual abstraction probed, rather than reflecting a uniform loss across the visual hierarchy. Future studies may consider a more robust approach examining representational shifts on a continuum (see Davis et al., 2021).

Our visual vulnerability results directly link declines in visual representational strength to age-related reductions in structural connectivity. These findings extend previous research showing white matter integrity has broad implications for functional brain data and highlight the critical role of brain architecture in shaping functional and behavioral aging effects (Madden et al., 2020), particularly in the study of working and episodic memory (Bennett & Rypma, 2013; Davis et al., 2009, 2018; Lockhart et al., 2012). Our mediation analyses suggest that declines in visual representation are statistically accounted for by age-related reductions in white matter integrity linking visual representation regions. However, white matter integrity is unlikely to be the only structural factor relevant to VSRS. Investigations on the roles of gray matter volume and brain iron accumulation (Howard et al., 2022) in the VSRS phenomenon are necessary next-steps. Additionally, these analyses are cross-sectional and mediation effects should be interpreted as statistical dependencies rather than evidence of causal direction, and future research should probe the structural underpinnings of age-related representational changes longitudinally.

Our findings regarding semantic resilience show that older adults showed stronger semantic representational strength in semantic representation regions. Coupled with the age-related declines in visual representation, we interpret our second finding as evidence of semantic resilience (the capacity of semantic systems to remain available, and in some contexts become more behaviorally relevant, when visual representations are weakened). This semantic resilience refines our understanding of prior VSRS findings (Deng et al., 2021; Naspi et al., 2023) as well as converging evidence that mnemonic semantic representations are preserved, and in some cases sharpened, in older adults, whereas contextual and more detailed information is more vulnerable representational decline (Ehrlich et al., 2024). We note that in our data, this resilience was not uniform across individuals. Instead, it depended on preserved connectivity within semantic representation regions. Additionally, the absence of a mediating effect of semantic structural connectivity on the age-semantic representation is theoretically informative. Unlike visual representational decline, which appears to track age-related structural degradation more directly, increased semantic representation may not be a simple consequence of aging or of connectivity changes alone. Instead, semantic resilience may be conditional: preserved structural connectivity of semantic regions may create the capacity for semantic recruitment, but whether that capacity is expressed likely depends on task demands, available semantic knowledge, and the degree of visual-system compromise. This interpretation is consistent with our moderation findings, which showed that connectivity within semantic representation regions predicted semantic representational strength specifically in older adults.

Our moderation analyses provided critical linkages between modalities that help explain the role of semantic representations within the VSRS phenomenon and strong evidence that semantic representations can serve a compensatory function in aging. Compensation is most convincing when a neural resource supports behavior specifically under conditions of deficit. That is the pattern observed here (Cabeza et al., 2018). Among older adults, semantic representations were most strongly related to perceptual memory when visual-system structural connectivity was low. Thus, semantic resilience did not reflect a general age-related increase in abstraction, but a conditional mechanism that became behaviorally relevant when vulnerability in visual circuitry was greatest. Further, this finding refines our understanding of brain-behavioral resilience in VSRS as it clarifies that successful resilience depends not only on preserved knowledge, but also on preserved pathways that allow that knowledge to be deployed. Future work should investigate whether semantic support mechanisms can be leveraged to promote resilience and consider the intraindividual variability in neural plasticity in the face of brain aging (J. Merenstein et al., 2024).

To conclude, our findings suggest that age-related representational change reflects both decline and adaptation. Structural and visual representational losses hamper memory performance, but these effects are partly offset by reliance on the semantic systems that provide compensatory support. This balance illustrates how the aging brain reorganizes its resources, relying on preserved pathways to sustain performance in the face of decline. Appreciating this dual pattern of vulnerability and resilience not only deepens our understanding of neural aging, but also suggests that interventions targeting semantic processing may help amplify compensatory mechanisms that preserve memory across the lifespan.

## Online Methods

### Participants

Thirty-eight younger adult (YA, ages 18-30) and 38 older adult (OA, ages 60-85) participated in the study and were monetarily compensated. Eligibility requirements included: ages between 18 and 30 years or between 60 and 85 years, English fluency, no history of significant neurological or psychiatric conditions, and not currently taking medications that affect cognition or cerebral blood flow except for antihypertensive agents. OAs were also screened for cognitive impairment with the Montreal Cognitive Assessment (MoCA) and obtained a minimum score of 23 (Hobson, 2015). Participants provided written informed consent and the study protocol was approved by the Duke University Health System Institutional Review Board and all methods were performed in accordance with the relevant guidelines and regulations. Four YAs and eight OAs withdrew from the fMRI study, and one additional YA was excluded due to failure to provide task responses, resulting in final fMRI samples of 33 YAs and 30 OAs. All of these participants were included in fMRI analyses. Due to technical issues and time constraints, three YAs and four OAs did not undergo diffusion-weighted imaging, yielding final samples of 30 YAs and 26 OAs with both fMRI and DTI data.

### Behavioral methods

#### Paradigm

The study took place in three sessions over 2 days. This sample size aligns with other studies probing age-related changes in representation (Deng et al., 2021; Hill et al., 2021; Naspi et al., 2023; Srokova et al., 2024). An expansive outline of the study design for the tasks described below can be found in (Huang et al., 2024). Here, we focus on the Object Perception and Memory Retrieval tasks.

During all sessions, participants underwent fMRI scanning at Duke University’s Brain Imaging Analysis Center while performing perception and memory tasks. The first day (**Figure 1A**) consisted of diffusion-weighted imaging (DWI), and fMRI scanning of the object encoding phase. The object perception phase consisted presentations of a series of 114 objects with corresponding labels during 3 runs in an fMRI scanner (38 trials per run). Participants were asked to rate how well the words describe the object on a 4-point scale. The objects were presented for 4 seconds, followed by a jittered fixation cross with an average duration of 4 seconds.

**Figure 1:**
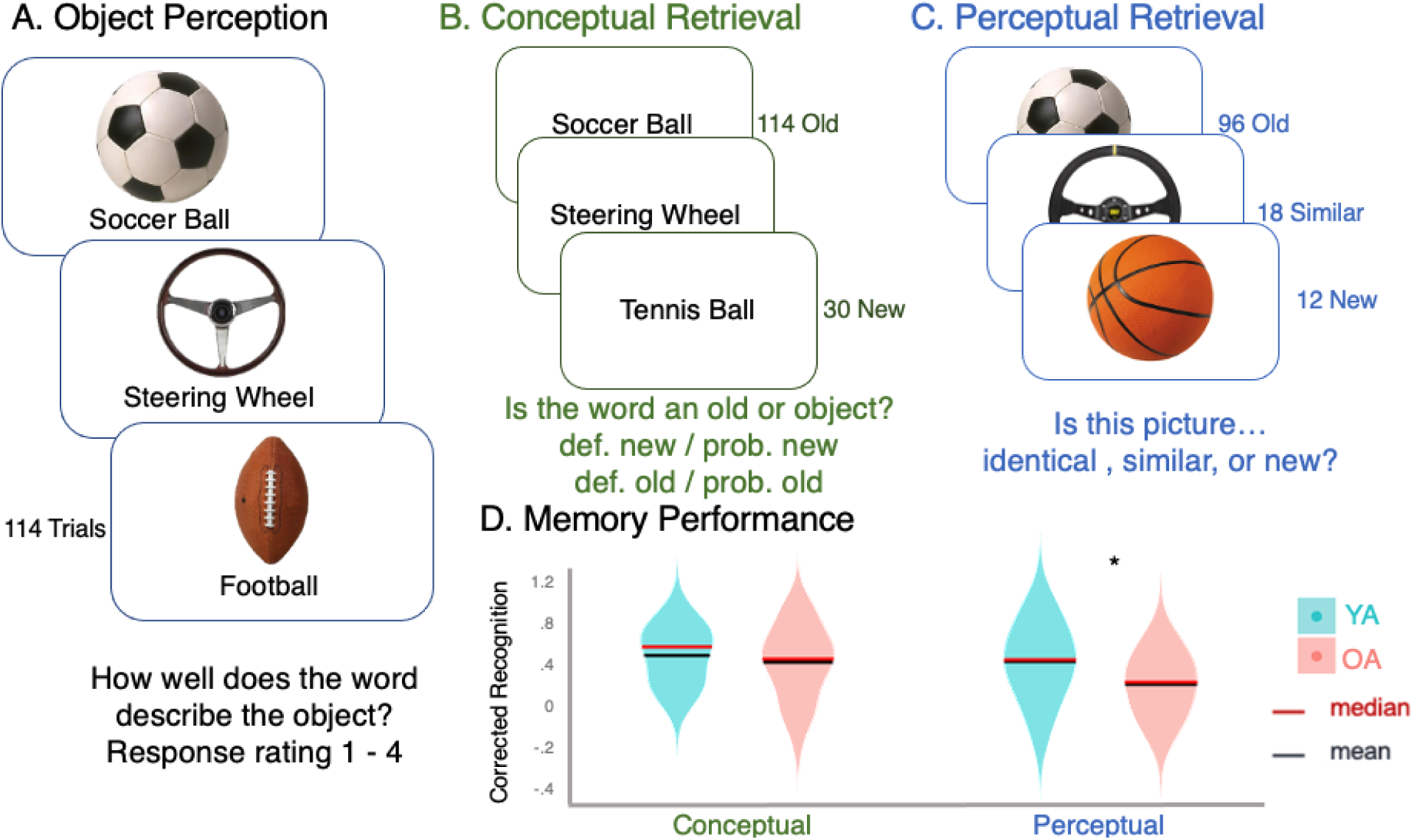
Experimental Paradigm and Behavioral Performance. A) While undergoing fMRI scanning, participants view 114 objects and object labels while judging the extent the label describes the object. B. Participants are tested for their conceptual memory of the objects and indicate their confidence on each trial. C. Participants are tested for their perceptual memory of the objects, indicating if each image is “old”, “Similar”, or “New”. D. Violin plots of Corrected Recognition scores during conceptual (green) and perceptual (blue) memory retrieval. Younger adults (YAs) in teal, older adults (OAs) in salmon. * Indicates significant two-sample t-test at p < 0.05.

A Memory Encoding session took place on the second day of testing, at least 7 days after Object Perception. During Encoding, participants viewed images of 114 unique real-world scenes along with the 114 objects they had seen in during Object Perception. In each trial, participants were shown a scene image for 3 seconds, followed by a jittered empty box indicating the continuation of the trial, which lasted for an average of 3 seconds, and finally an object image for 4 seconds. During the object presentation, participants rated “how likely it is to find the object in the scene” on a 4-point scale. Each trial was separated by a jittered fixation cross with an average duration of 4 seconds.

Twenty-four hours later, the final session on the third day consisted of two retrieval tasks: conceptual (**Figure 1B**) and perceptual (**Figure 1C**). During the conceptual retrieval phase, participants were shown 114 words previously shown in both encoding phases (old trials) and 30 new words. Participants were asked to indicate whether each word was shown previously and their confidence judgment on a 4-point scale (definitely new, probably new, probably old, definitely old). During the perceptual retrieval phase, participants indicated if the object shown is identical to one shown during the encoding phases (96 “old” trials), the same concept but a different visual exemplar (18 “similar” trials), or not shown during the encoding phases (12 “new” trials).

#### Behavioral analyses

Memory performance was evaluated using the perceptual memory test. We excluded ‘new’ trials, as well as trials in which participants responded ‘new,’ since these reflect tests of conceptual memory or conceptual errors, respectively. The hit and false alarm rates were calculated, and by subtracting the latter from the former, derived a corrected recognition measure (CR). We focused on corrected recognition because it accounts for response biases, such as frequently guessing “old,” which can artificially inflate hit rates. By subtracting the false alarm rate from the hit rate, this measure provides a more accurate assessment of memory sensitivity, the participant’s ability to distinguish between old and new items. Next, OA and YA perceptual and conceptual CR scores were submitted to a two-sample t-test to detect age differences in behavior.

### Neuroimaging methods

#### Structural & Functional MRI

MRI data were collected using a 3T GE MR750 Scanner equipped with an 8-channel head coil at the Brain Imaging and Analysis Center (BIAC) at Duke University. Each MRI session began with a localizer scan, during which 3-plane (straight axial/coronal/sagittal) faster spin echo images were obtained. A high-resolution T1-weighted (T1w) structural scan was then acquired, consisting of 96 axial slices parallel to the AC-PC plane, with voxel dimensions of 0.9 × 0.9 × 1.9 mm³. This was followed by blood-oxygenation-level-dependent (BOLD) functional scans using a whole-brain gradient-echo echo planar imaging sequence (repetition time = 2000 ms, echo time = 30 ms, field of view = 192 mm, 36 oblique slices with voxel dimensions of 3 x 3 x 3 mm³). Task instructions and stimuli were delivered using the PsychToolbox program (Kleiner et al., 2007) and projected onto a mirror at the back of the scanner bore. Participants responded using a four-button fiber-optic response box. To reduce scanner noise, participants wore earplugs, and MRI-compatible lenses were provided when necessary to correct vision. Foam padding was placed inside the head coil to minimize head movement.

#### Diffusion weighted imaging

On the first day of MRI scanning, participants underwent whole-brain diffusion-weighted imaging (DWI) with a multiplexed sensitivity-encoding (MUSE) sequence, which allows for high spatial resolution (1mm3 isotropic voxels), improved signal to noise ratio, and decreased motion artifacts (Chen et al., 2013; J. L. Merenstein et al., 2023). Single-band data were acquired using a spin-echo EPI sequence with these parameters: voxel size = 1 × 1 × 1 mm, acquisition matrix = 256 × 256 × 90 mm, TR = 13 000 ms, TE = 58 ms, flip angle = 90°, SENSE factor = 1 and multi-band factor =1. Diffusion-weighted gradients were applied in 15 directions with a b value of 1000 s/mm2 and with two non-diffusion-weighted b = 0 s/mm2 images.

#### Trial-Level Representational Similarity Analyses (tRSA)

Our analytical approach for representational strength utilized trial-level representational similarity analysis (tRSA) methods (**Figure 2A)** (Huang* & Howard* et al., 2025). First, two model RSMs were constructed to capture visual and semantic similarity across the 114 objects. Importantly, our goal was not to adjudicate among competing models of visual or semantic similarity, but to test whether age-related representational changes align with independently defined visual versus semantic similarity structures. For the visual model RSM, we aimed to model robust mid-stream visual similarity which we derived from C2 features in the Hierarchical Model of Object Recognition (HMAX), implemented in PsyTorch. The C2 layer provides a compact feature vector that reflects complex visual properties (e.g., shape/configuration) with invariance to position and scale. (Kriegeskorte et al., 2008; Riesenhuber & Poggio, 1999, 2002; Serre et al., 2005, 2007; Sufikarimi & Mohammadi, 2020). Pairwise similarity between objects was computed using Pearson’s r to generate a 114 × 114 visual RSM. For the semantic model RSM, semantic similarity was defined using word vector representations based on corpus-level co-occurrence statistics (Mikolov et al., 2013), and cosine distance was used to build the corresponding 114 × 114 semantic RSM.

**Figure 2:**
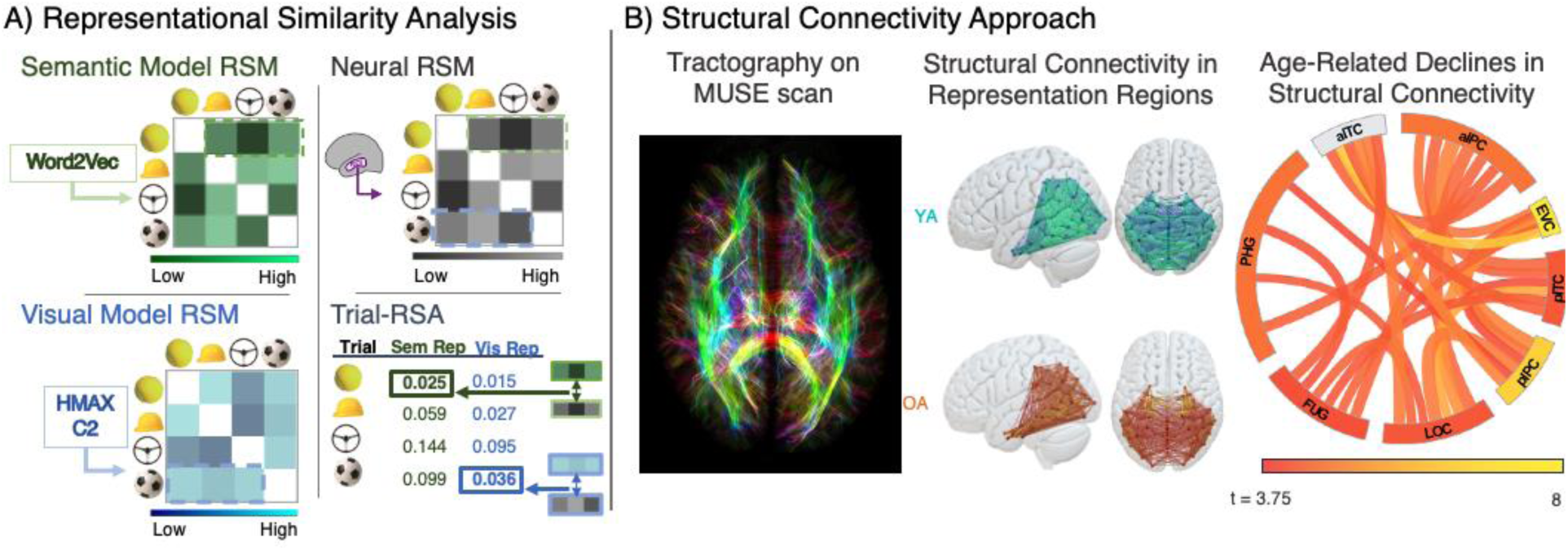
**Analytical Approaches**. *A) Representational Similarity Analysis. Representational* similarity matrices were constructed for visual (blue) and semantic (green) representation types. Participant-specific neural Representational Similarity Matrices (RSMs) (gray) were constructed for each region in the Brainnetome atlas. Each item’s unique neural similarity vector is correlated with that item’s visual and semantic representational similarity vector, yielding a visual and semantic trial-level representational similarity value, respectively. *B) Structural Connectivity Approach.* The high-resolution multiplexed sensitivity-encoding (MUSE) diffusion imaging scan was used to detect tracts across the whole brain. Next, the average fractional anisotropy along detected streamlines connecting each region of the Brainnetome Atlas was used to construct the structural connectivity matrices. Next, the connectomes were filtered to only include edges connecting the 8 broad representation regions of interest. The mean connectomes are visualized separately for younger (teal) and older (orange) adults. Finally, age group two sample t-tests were performed on each edge connecting representation regions. Significant (fdr corrected p<.05) t-values (red to yellow scale) were then averaged over the 8 broad regions of interest. The color of the region labels indicates the strength of the age effect among edges within the same broad region of interest on the same red to yellow color scale.

Next, we obtained whole-brain single-trial beta estimates from each session for each participant (Mumford et al., 2012). We then utilized our internal RSA toolbox to obtain activity patterns for each subregion in the Brainnetome atlas (Fan et al., 2016) that overlap with eight broad regions of interest: the early visual cortex (EVC), lateral occipital cortex (LOC), fusiform gyrus (FUG), parahippocampal gyrus (PHG), anterior part of the inferior temporal cortex (aITC), posterior part of the inferior temporal cortex (pITC), anterior part of the inferior parietal cortex (aIPC) and the posterior part of the inferior parietal cortex (pIPC). We then constructed participant-specific single-trial estimates during object perception and constructed neural RSMs for each sub region. The cells of the neural RSM represented the similarity of voxel patterns (Pearson correlation) in fMRI activation patterns by stimuli. Then, we computed trial-level representation values by utilizing tRSA. tRSA values represented the correlation of stimulus properties in the RSMs and the fMRI activity patterns in the neural RSM for each object (the vector on each row in the matrices) and are an index of a region’s representational strength during a trial (Davis et al., 2021; Huang et al., 2024; Huang* & Howard* et al., preprint; Yu et al., 2025). This approach is a variation of traditional representational similarity analysis approaches in which second-order correlations between neural RSM and model RSMs are computed using the entire matrix (Kriegeskorte & Kievit, 2013).

Next, tRSA values were submitted to a series of mixed-effect linear models to identify representation regions that exhibit an age-related change. In each broad region of interest, a model containing the interaction of Age Group (older adults and younger adults) and Representation Type (Semantic and Visual) and their main effects, as well as the random intercepts of Participant, Stimuli, and Sub-Region of Interest was constructed. Hypothesized representation regions were classified depending on the significance and nature of the Age Group by Representation Type interactions. Any region exhibiting an Age Group by Representation Type interaction will be further classified into age-related visual or semantic representational change regions if the tests reveal the hypothesized age effect(s) in the region contributed to the interaction.

#### Structural connectivity

Raw diffusion-weighted imaging (DWI) data were processed using MRtrix3 (Tournier et al., 2019). Data were denoised (dwidenoise), corrected for motion and eddy current–induced distortions (dwifslpreproc), bias-field corrected (dwibiascorrect), and skull-stripped to generate whole-brain masks (dwi2mask). All images were visually inspected and deemed acceptable with respect to mask coverage, distortion correction, and absence of imaging artifacts.

Structural connectivity was assessed by estimating fiber orientation distribution (FOD) maps using multi-tissue constrained spherical deconvolution (CSD; Tournier et al., 2019). Tissue-specific response functions were derived using the dhollander algorithm, with three compartments (GM, WM, CSF). A gray-matter/white-matter (GM/WM) boundary mask was generated from the T1-weighted image (5ttgen) and registered to native diffusion space.

Anatomically constrained probabilistic tractography was performed in native diffusion space, with streamlines seeded from and terminated within the GM/WM boundary mask (Smith et al., 2012). Tractography parameters included a minimum streamline length of 1 mm, a maximum length of 250 mm, and an FOD cutoff of 0.06. For each participant, 10 million streamlines were generated (tckgen) and weighted using spherical-deconvolution informed filtering (SIFT2; tcksift2) to reflect apparent fiber density (Smith et al., 2015).

Fractional anisotropy (FA) values were sampled along streamlines, and the mean FA for streamlines connecting each pair of regions was used to weight connection strength, yielding FA-weighted structural connectivity estimates that integrate macrostructural and microstructural properties. Streamlines were transformed to MNI152 1 × 1 × 1 mm space (tcktransform), and symmetric 246 × 246 structural connectivity matrices were constructed using tck2connectome based on the Brainnetome Atlas (Fan et al., 2016). Streamline endpoints were assigned using a 2 mm radial search, and connection weights were normalized for regional volume.

#### Principal axis factor analysis

We conducted a principal axis factoring (PAF) on the relevant indicator variables (structural connectivity, visual representation strength, and semantic representation strength, for all participants) and used the factor score for each participant from the first unrotated factor as the summary measure. We used a PAF rather than another factor approach because our interest was in the shared variance among the indicator variables rather than the mathematically independent components (Salthouse et al., 2015), and this method has been used in a variety of cognitive aging studies (Howard et al., 2022; Madden et al., 2020). PAF was conducted using the fa() function from the *psych* package in R (Revelle, 2015). This method was chosen to capture the shared variance from the observed variables while accounting for measurement error. The factor analysis was performed by specifying the correlation matrix of the subset as the input, extracting one factor, and selecting the principal axis factoring method. The output included standardized factor loadings, communalities (the proportion of variance in each variable explained by the factor), unique variances (the variance not explained by the factor), and the proportion of total variance accounted for by the extracted factor. This approach allowed for a robust exploration of the relationships among the observed variables and exclusion of ROIs that did not contribute sufficiently to the principal factor (pa < .20). Model adequacy was evaluated using indices of factor score adequacy, which assess how well the estimated factor scores represent the latent construct. Following conventional guidelines (Gorsuch, 2014; MacCallum et al., 1999).

Four PAFs were conducted using data from visual and semantic representational change regions, which were identified in a data-driven manner depending on the significant age-group effects in the representational mixed effects models. The first PAF was *visual representational strength*. To create this factor, ROIs with significant age-related differences in visual representational strength were selected. For those ROIs, the model estimate of visual representational strength for each participant was extracted from the representational mixed effect model described above. The second PAF, *structural connectivity of visual representation regions* to all regions within the representation network was calculated. Similarly, third PAF *semantic representational strength* was created with data from ROIs that exhibited age-related changes in semantic representation strength, and the final PAF, *structural connectivity of semantic representation regions* to all regions within the representation network was calculated.

### Mediation and moderation analysis

Our multimodal data consisting of structural connectivity of representation regions, semantic knowledge, visual and semantic representational strength, perceptual memory performance, and participant age group was submitted to a series of moderation and mediation models in R with the lavaan package (Rosseel, 2012). Our first set of mediation and moderation models assessed the relationships of: age group on structural connectivity (*a* path), structural connectivity on visual and semantic representational strength (*b* path), age group on visual and semantic representational strength (total, *c* path), and finally, the extent the age and representational relationships are mediated by structural connectivity (indirect path, *a*b*). Our second mediation and moderation models assessed the drivers of older adult memory performance by analyzing the relationships of: visual and semantic representational strength structural connectivity (*a* path) structural connectivity and semantic knowledge on visual memory performance (*b* path), visual and semantic representational strength on conceptual and perceptual memory performance (total, *c* path), and finally, the extent the representational strength and memory performance relationships are mediated by structural connectivity and semantic knowledge (indirect path *a*b*). Data from younger adults were submitted to the same models to confirm the effects are specific to the older adult sample.

## Acknowledgements

We thank Lamont Conyers and Jennifer Graves for their extensive MRI support and all participants for their participation.

## Funding

This study was supported by the National Institutes of Health, grant numbers: R01AG066901 and R01AG075417.

## Author Contributions

Cortney M. Howard: Conceptualization; Formal analysis; Writing—Complete 1st draft; Methodology. Shenyang Huang: Formal analysis; Methodology. Kirsten Gillette: Data acquisition; Data quality assurance. Lifu Deng: Conceptualization; Data acquisition. Roberto Cabeza: Conceptualization; Funding acquisition; Methodology; Supervision; Writing—Review & editing. Simon W. Davis: Conceptualization; Formal analysis; Funding acquisition; Methodology; Supervision; Writing—Review & editing.

## Notes

### Competing Interest Statement

The authors have declared no competing interest.

